# Subthreshold Kir and I_h_ currents modulate excitability of Layer 1 VIP interneurons in the medial prefrontal cortex

**DOI:** 10.64898/2026.01.28.702118

**Authors:** Claudio Moreno, Denise Riquelme, Camila Cornejo, Paula Leyton, Elias Leiva-Salcedo

## Abstract

Cortical layer 1 (L1) is a key site for integrating top-down and bottom-up information and is populated by inhibitory interneurons, including vasoactive intestinal peptide (VIP)-expressing cells. These interneurons regulate information flow across the cortical column by disinhibiting pyramidal neurons, yet the subthreshold ionic mechanisms that shape their excitability in the medial prefrontal cortex (mPFC) remain poorly understood. Here, we characterize the electrophysiological properties of L1b VIP interneurons in mouse mPFC, focusing on the role of the hyperpolarization-activated cation current (I_h_) and inward-rectifying potassium current (Kir). Using whole-cell recordings in acute slices, we find that L1b VIP interneurons exhibit constitutively active I_h_ and Kir conductances. Inhibition of I_h_ with ZD-7288 hyperpolarized the resting membrane potential (RMP), increased input resistance (R_in_), and prolonged action potential (AP) duration without altering rheobase or firing frequency, while increasing EPSP-spike coupling probability. Blocking Kir with BaCl_2_ depolarized the RMP, increased R_in_ and membrane time constant, reduced rheobase, increased firing frequency, and similarly enhanced EPSP-spike probability and AP duration. Voltage-clamp experiments confirmed the presence of ZD-7288-sensitive I_h_ and BaCl_2_-sensitive Kir currents of small amplitude but operating on a high-resistance membrane, consistent with a strong impact on excitability. In contrast, neither I_h_ nor Kir inhibition affected the amplitude or frequency of spontaneous or miniature EPSCs, indicating that these currents do not measurably alter basal excitatory synaptic transmission. Immunofluorescence revealed weak somatic HCN1 and no detectable HCN2 expression in L1b VIP interneurons. Notably, Kir inhibition unmasked an I_h_-dependent voltage sag during hyperpolarizing steps, suggesting that constitutive Kir activity normally masks I_h_ recruitment at subthreshold potentials. Together, these results indicate that I_h_ and Kir are active near RMP in L1b VIP interneurons and jointly regulate their passive properties, intrinsic excitability, and EPSP-spike coupling, therefore shaping how L1 VIP cells filter incoming signals and influence information flow within the prefrontal cortical column.

## 1. INTRODUCTION

Layer 1 (L1) of the neocortex is a critical site for the convergence of long-range cortical, thalamic, and neuromodulatory inputs [1]. In the medial prefrontal cortex (mPFC), this convergence is central for coordinating attention, working memory, and executive functions through the integration of top-down and bottom-up information streams [2]. Unlike deeper layers, L1 contains no excitatory somata and is composed exclusively of inhibitory interneurons, primarily LAMP5 and VIP cells, [3]. LAMP5 neurogliaform cells are located in the upper part of L1 (L1a) and mediate widespread GABAergic inhibition of pyramidal apical dendrites [4–6]. Meanwhile VIP interneurons are located in the bottom part of L1 (L1b) and disinhibit pyramidal neurons by inhibiting Somatostatin (SST) and Parvalbumin (PV) expressing interneurons, therefore enhancing the impact of top-down signals [7–9]. This positions VIP interneurons as key modulators of cortical column excitability and state-dependent information routing.

VIP interneurons in L1 and at the L1-L2 border display two principal morphologies: a bipolar form with vertically oriented dendrites extending through Layers 1 and 2, and a multipolar form with horizontally ramified dendrites. Both morphological types are positioned to sample dense neuromodulatory inputs and long-range corticocortical projections [2,10]. Their axons primarily target layers 2/3 and 5, enabling them to coordinate disinhibition across multiple layers. Although their anatomical organization in sensory cortices has been described [11–13], their intrinsic electrophysiological properties, particularly in mPFC L1b, remain poorly understood. Most functional studies focus on VIP cells in deeper layers, where they are characterized by high intrinsic excitability and robust disinhibition [14]. Yet the ionic mechanisms that shape their excitability in L1, a region dominated by distal and modulatory inputs, have received little attention.

Subthreshold ionic conductances are particularly relevant because they determine how neurons integrate synaptic inputs below spike threshold, influencing firing probability, temporal precision, and excitability dynamics [15–19]. Among these, the inward-rectifying potassium current (Kir) and the hyperpolarization-activated cation current I_h_ may play an especially important role in modulating the function of L1 interneurons, due to their strong filtering and gain-control capabilities [16,20]. Inward rectifying potassium channels (Kir) stabilize the resting membrane potential, decrease membrane resistance and slow its responses to voltage changes while also promoting an influx of potassium at hyperpolarized membrane potentials, acting as one of the main filters for weak synaptic inputs [21,22]. By contrast to Kir, the hyperpolarization-activated cation current I_h_ depolarizes the membrane, accelerates membrane responses, and promotes coincidence detection while contributing to resting potential instability through activity-dependent inactivation [23,24]. Although Kir and I_h_ have been well characterized in pyramidal neurons, Parvalbumin-expressing and Somatostatin-expressing interneurons, the expression, strength, and functional consequences of these currents in VIP interneurons remain largely unknown, despite suggestive electrophysiological evidence of their presence in this interneuron [12,25].

Understanding these mechanisms is particularly critical in L1, where VIP interneurons may act as gatekeepers of cortical disinhibition and, consequently, of top-down information flow. In contrast to LAMP5 neurogliaform cells, which are abundant and provide widespread inhibition [4–6,26–28], L1 VIP interneurons constitute only ∼10% of L1 cells and remain understudied despite their strategic position and powerful disinhibitory function. In the mPFC, VIP interneurons are located predominantly in the lower part of layer 1 (L1b) and at the interface between L1b and layer 2 [2,3,29,30]. Their dendritic access to neuromodulatory and top-down signals, combined with axonal outputs to layers 2/3 and 5 [10], make them ideally suited to regulate the balance between contextual information and sensory input [7,8,31,32]. Subthreshold currents such as Kir and I_h_, by shaping resting membrane potential, input resistance, and EPSP-spike coupling, are therefore likely to be fundamental determinants of how VIP cells filter incoming signals and influence circuit computations.

However, the combinatorial operation of Kir and I_h_ in L1 VIP interneurons, and its impact on neuronal excitability and spike-timing, has not been systematically examined. Here we hypothesize that the balance of I_h_ and Kir conductances in VIP interneurons of mPFC L1b is a major determinant of their excitability and spike-timing, therefore shaping their disinhibitory influence across cortical layers. To test this hypothesis, we combined whole-cell patch-clamp recordings with immunofluorescence to characterize Kir and I_h_ in L1 VIP cells. We find that both conductances are constitutively active and strongly regulate passive membrane properties, synaptic integration, and firing precision. These results reveal that Kir and I_h_ fine-tune VIP interneuron excitability and suggest a modulatory role in the flow of top-down and bottom-up information through the cortical column with implications for sensory processing flexibility, network stability, and cognitive control.

## 2. METHODS

### 2.1. Animals

Male mice between 30 and 90 days old were used for the study (median = 56). For experiments, we crossed homozygous STOCK Vip^tm1(cre)Zjh/J^ male mice with B6.Cg-Gt(ROSA)26Sor^tm9(CAG-tdTomato)Hze/J^ homozygous female mice. All mice were maintained in a 12-hour light-dark cycle, with the light cycle occurring between 7:00 AM and 7:00 PM. Animals had *ad libitum* access to water and food. All experiments were conducted following the animal protocol (N°: 193/2022) approved by the Ethical Committee of the Universidad de Santiago de Chile according to the rules and guidelines of the National Agency of Research and Development (ANID-Chile).

### 2.2. Electrophysiological recordings

#### Slice preparation

300 µm acute brain slices were prepared from mice deeply anesthetized with isoflurane (3%) and intracardially perfused with ice-cold cutting solution containing: 125 NaCl, 2.5 KCl, 5 MgCl_2_, 0.5 CaCl_2_, 1.25 NaH_2_PO_4_, 25 NaHCO_3_, 11 Glucose (pH 7.4). Following dissection, tissue blocks containing the prefrontal cortex were cut using a vibratome. Then, slices were transferred to a recording chamber with oxygenated ACSF containing: 125 NaCl, 2.5 KCl, 1.3 MgCl_2_, 2.5 CaCl_2_, 1.25 NaH_2_PO_4_, 25 NaHCO_3_, 11 Glucose (pH 7.4, ∼300 mOsm/kg). After 1 h recovery, the slices were transferred to a recording chamber mounted on a Zeiss Axio Examiner A1 DIC microscope. Slices were continuously perfused with oxygenated ACSF (2-3 mL/min) at 34 ± 2 °C.

#### Patch clamp

Whole-cell recordings were performed in layer 1 of the mPFC in tdTomato expressing VIP interneurons, using borosilicate glass pipettes (4 to 6 MΩ). For voltage and current clamp, the intracellular solution contained (in mM): 120 K-gluconate, 10 KCl, 8 NaCl, 10 HEPES, 0.5 EGTA, 4 Mg-ATP, 0.3 Na-GTP, pH 7.2 adjusted with KOH (∼300 mOsm/kg). For voltage clamp recordings, pipette, and whole-cell capacitance were compensated, series resistance was continuously monitored and compensated by 80%; neurons were held at -80 mV. The voltage-step protocols consisted in voltage steps from -130 to -50 mV with 10 mV steps from a holding potential of -80 mV, with a duration of 0.5 s and delivered somatically at 0.2 Hz. Liquid junction potential was 16.8 mV and was not subtracted (calculated according to the stationary Nernst-Planck equation using LJPcalc software (https://swharden.com/LJPcalc)).

The current-step protocols consisted of a series of somatic current steps from -60 to 200 pA in steps of 20 pA with a duration of 1 s and delivered at 0.2 Hz at a holding current of 0. All current-clamp recordings were bridge-balanced and pipette capacitance neutralization was applied. Recordings were monitored continuously for changes in R_s_ using a pulse of 50 and -50 pA at the end of each current step. Recording showing changes in R_s_ > 20% were discarded from the analysis. Voltage and current clamp recordings were performed using a Multiclamp 700A (Molecular Devices, USA), digitized with a National Instruments PCIe-6323. The signal was low-pass filtered at 10 kHz and digitized at 50 kHz using WinWCP 5.8 (https://github.com/johndempster/WinWCPXE/releases/tag/V5.7.8).

#### Evoked EPSPs

To perform the extracellular stimulation protocol, a tungsten bipolar electrode (FHC) was positioned in layer 1 (L1) of the mPFC, approximately 200 µm from the soma of the recorded VIP interneuron. Stimuli were delivered every 3 s as 50 µs test pulses via an A365 stimulus isolator (WPI, USA). The stimulus intensity (range: 0.1-1.0 mA) was calibrated for each cell to elicit an action potential (AP) with a probability of 0.5 (determined over 20 traces). To isolate the excitatory component, all experiments were conducted in the presence of 50 µM picrotoxin unless otherwise noted. For voltage-clamp experiments isolating subthreshold I_h_- and Kir-mediated currents, the solution was supplemented with 5 mM 4-AP, 5 mM TEA, and 0.5 µM TTX. I_h_ was inhibited using 20 µM ZD-7288 (Tocris, USA), and Kir was inhibited using 50 µM BaCl_2_ (Sigma-Aldrich, USA).

### 2.3. Tissue immunofluorescence

Coronal brain slices containing the prefrontal cortex were prepared from mice aged 30 to 90 days. Briefly, male mice were deeply anesthetized with 3% isoflurane. Animals were intracardially perfused with 0.1 M PBS (pH 7.2) followed by freshly made 4% w/v formaldehyde dissolved in 0.1 M PBS (pH 7.4). All mice were euthanized by decapitation, and brains were quickly removed and placed in 4% w/v formaldehyde (Merck, Germany) for overnight incubation at 4°C. The next day, brains were placed in a vibrating tissue slicer and sectioned using a sapphire blade to obtain 60 µm thick slices. Slices were obtained between the anteroposterior coordinates 1.54 to 2.46 mm from the bregma.

#### 2.3.1. Antibodies

Monoclonal anti-HCN1 (N70/28, RRID: AB_2750810) and anti-HCN2 (N71/37, RRID: AB_2279449) antibodies were obtained from NeuroMab (United States). Polyclonal Anti-MAP2 (RRID: AB_776174) antibody was obtained from Abcam (United States). Secondary Alexa Fluor 488 Goat anti-mouse IgG (H+L) (A-11029), and Alexa Fluor 546 donkey anti-rabbit IgG (A-10040) were obtained from Thermo Fisher Scientific (United States).

#### 2.3.2. Immunofluorescence Labeling

Tissue sections containing the prefrontal cortex were permeabilized in 0.3 % Triton X-100 (TX-100) for 10 min and then blocked in 0.3% BSA, 0.09 % TX-100 and 7% normal goat serum (NGS) in 0.1 M PBS (pH 7.4) for 2 hours at room temperature (RT). Sections were then incubated with the primary antibody: HCN1 (1:50), HCN2 (1:100) or MAP2 (1:300) diluted in 1% BSA plus 0.3 % TX-100 with gentle rocking for 24 hours at 4°C. For negative controls, tissue sections were incubated with blocking solution and without the primary antibody. Sections were washed in 0.1 M PBS for 30 minutes before incubation with the secondary antibody (1:500 dilution, 1.5 hours at RT). After six, 5-minute washes with PBS, the sections were mounted on glass slides using Mowiol mounting media and covered with a 0.17 mm thick coverslip. Fluorescence was observed in an LSM 800 inverted confocal microscope (Zeiss, Germany). High magnification confocal images were acquired using a 40x 1.1 N.A. water immersion objective, with a pinhole of 1 Airy unit for each channel, a scan zoom of 0.5, frame scan mode with averaging of 4, and a pixel time of 2.06 µs. All images were obtained at 8-bit pixel depth at 1024 x 1024 resolution. Images were taken as a Z-stack at 1 µm intervals and analyzed as Z-projections using Fiji-ImageJ. Images were adjusted for overall brightness, contrast, and level using Fiji-ImageJ.

### 2.4. Data analysis

#### Electrophysiological data analysis

Data were analyzed using Clampfit 10.3 and Igor Pro 6.37 with the Neuromatic module [33]. Rheobase was defined as the minimal current injection required to elicit an action potential (AP). The slope of the EPSP was calculated from 10-90% of the maximal amplitude. For EPSP-spike probability, an AP was considered when the slope of the rising phase of the AP reached 50 mV/ms and if the event passed a threshold of -10 mV. EPSP-spike latency was calculated by subtracting the time of EPSP onset (determined via the method described by [33]) from the time of spike threshold. Sag amplitude was calculated as the maximal hyperpolarization of the membrane potential between the first 100 ms after the onset of the current step minus the average values of the steady state of the last 500 ms of the hyperpolarization pulse. Spike precision was quantified as the coefficient of variation (CV) of EPSP-spike latency across trials.

#### Statistical analysis

Data are reported as the mean ± 95% confidence interval (C.I.), unless stated otherwise. Data normality was assessed using the Shapiro-Wilk test. Statistical significance to compare replicate means by row was evaluated using two-way ANOVA followed by Bonferroni post hoc tests. For multiple mean group analysis we used one-way ANOVA followed by a Tukey’s post hoc test. For parametric data paired *t*-tests were used in most of the two group analyses. For non-parametric paired data, the Wilcoxon matched pair signed rank test was used. Statistical significance was considered at *p*<0.05. All data analysis was performed using GraphPad Prism 9.

## 3. RESULTS

### 3.1. Localization and morphology of L1b VIP interneurons and lack of I_h_/Kir effects on spontaneous synaptic transmission

VIP interneurons in layer 1 (L1) are key integrators of bottom-up and top-down signals, and their excitability state strongly influences how these inputs are filtered and routed through the medial prefrontal cortex (mPFC). Subthreshold currents such as Kir and I_h_ play an important role in setting the resting membrane potential and shaping intrinsic responsiveness, providing neurons with strong filtering and gain-control properties. Therefore, characterizing these conductances is essential for understanding the computational role of L1 VIP interneurons. To investigate these mechanisms, we first examined the distribution and morphology of VIP interneurons in the mPFC of VIP^Cre^;Ai9 mice. Most labeled VIP somata were located in the lower subdivision of L1 (L1b), and no labeled somata were detected in the upper region (L1a) (Figure 1Ai). Morphologically, VIP cells were predominantly bipolar, with vertically oriented processes extending across L1 and L2, and in several cases projecting into the upper L1 compartment (Figure 1Aii).

**Figure 1.**
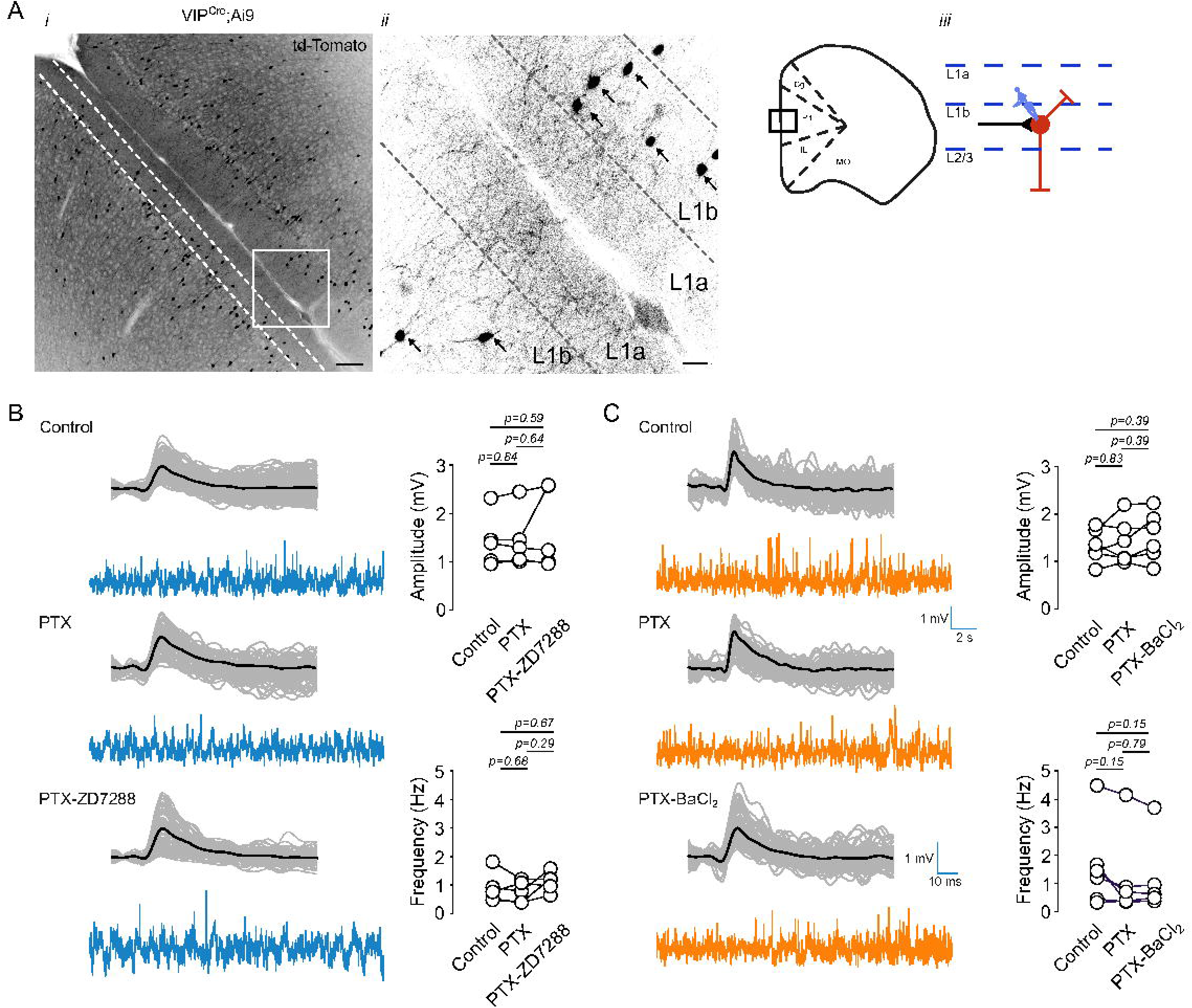
Contribution of Kir and I_h_ channels to sEPSPs in layer 1 VIP interneurons of the mPFC. (A) (i) Fluorescence image of the medial prefrontal cortex (mPFC) from VIP^Cre^;Ai9 mice expressing tdTomato. (ii) Higher magnification of the area indicated in (i); arrows indicate tdTomato-positive VIP interneurons in layer 1b. (iii) Schematic representation of the recording site. (B) Left: Representative sEPSP traces recorded in control conditions, in the presence of 50 µM picrotoxin (PTX), and with 50 µM PTX + 20 µM ZD-7288. Right: Summary plots quantifying the effect on sEPSP amplitude and frequency (n = 4 cells/4 animals). (C) Left: Representative sEPSP traces recorded in control, 50 µM PTX, and 50 µM PTX + 50 µM BaCl_2_. Right: Summary plots showing the effect on sEPSP amplitude and frequency (n = 4 cells/4 animals). Scale bars: 100 µm (Ai), 20 µm (Aii).

We next evaluated the contribution of I_h_ and Kir channels to spontaneous excitatory synaptic activity. In current clamp recordings performed in the presence of 50 µM picrotoxin, I_h_ inhibition with 20 µM ZD-7288 did not alter sEPSP amplitude or frequency (Figure 1B; Supplementary Table 1). Similarly, Kir inhibition with 50 µM BaCl_2_ produced no detectable change in sEPSP parameters (Figure 1C). To assess whether I_h_ or Kir influence quantal synaptic release or postsynaptic glutamatergic receptor responsiveness, we measured miniature EPSCs (mEPSCs) in voltage clamp in the presence of TTX and picrotoxin. Neither ZD-7288 (Figure 2A) nor BaCl_2_ (Figure 2B) affected mEPSC amplitude or frequency. These results demonstrate that I_h_ and Kir do not measurably influence basal excitatory synaptic transmission onto L1b VIP interneurons.

**Figure 2.**
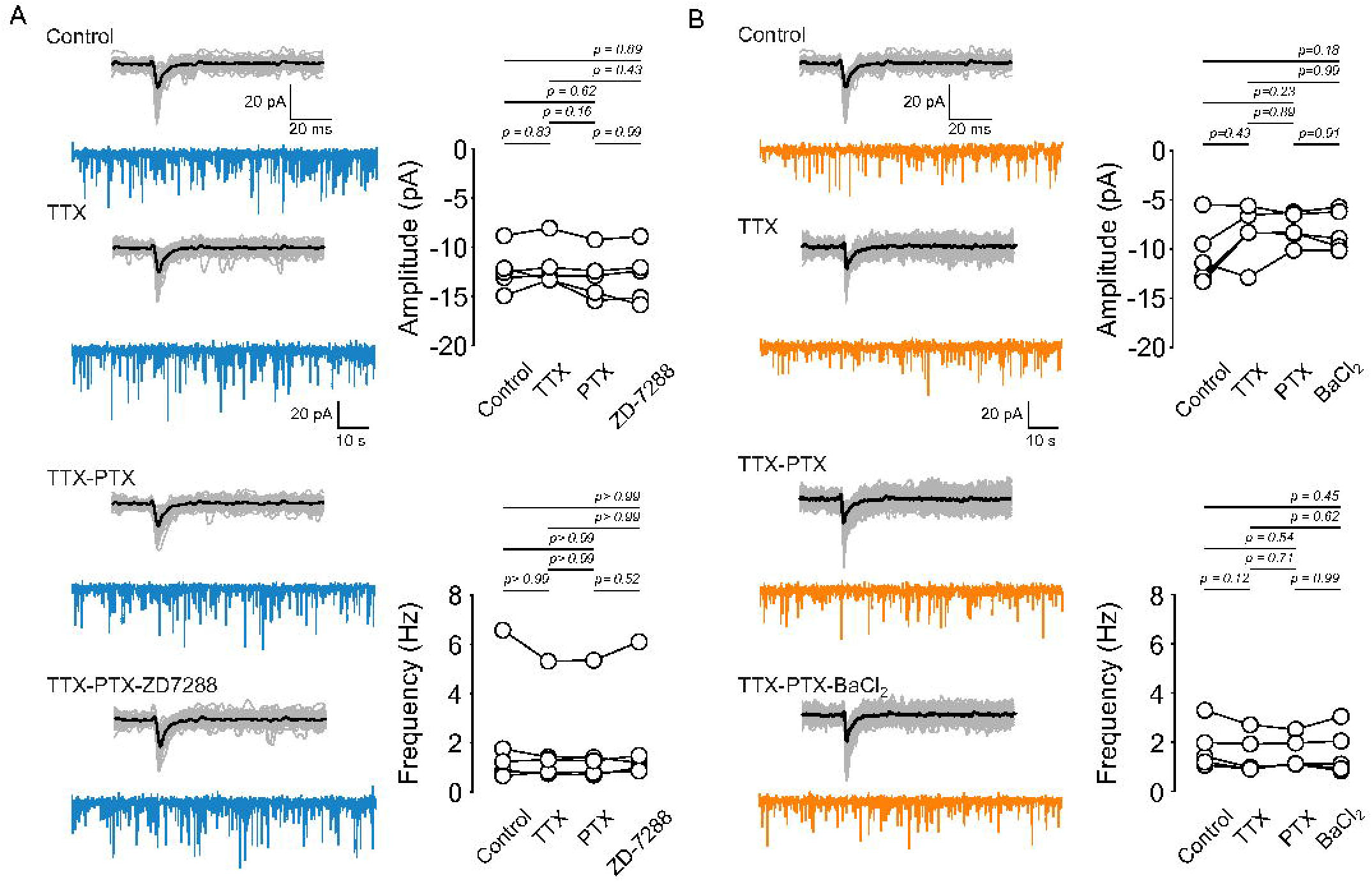
Effect of Kir and I_h_ inhibition on mEPSCs in layer 1 VIP interneurons of the mPFC. (A) Left: Representative current traces of spontaneous EPSCs and mEPSCs recorded in the presence of 0.5 µM TTX, 0.5 µM TTX + 50 µM PTX, and 0.5 µM TTX + 50 µM PTX + 50 µM ZD-7288. Right: Summary graphs showing the amplitude and frequency of synaptic currents. (B) Left: Representative current traces of spontaneous EPSCs and mEPSCs in the presence of 0.5 µM TTX, 0.5 µM TTX + 50 µM PTX, and 0.5 µM TTX + 50 µM PTX + 50 µM BaCl_2_. Right: Summary graphs showing the amplitude and frequency of synaptic currents.

### 3.2. Inhibition of I_h_ and Kir conductances increases EPSP-spike coupling in L1b VIP interneurons

To determine how I_h_ and Kir shape synaptic integration, we recorded EPSP-evoked spikes by stimulating excitatory afferents in L1b ∼200 µm from the soma (Figure 3A). Stimulation intensity was titrated for each neuron to achieve ∼0.5 action-potential probability under control conditions, enabling detection of bidirectional changes in synaptic responsiveness. Inhibition of I_h_ with 5 or 20 µM ZD-7288 increased spike probability to 1.0 in all recorded cells (Figure 3B, Supplementary Figure 3). Likewise, inhibition of Kir with 50 µM BaCl_2_ increased spike probability to 1.0 (Figure 3C). These effects persisted and were amplified when GABA-A receptors were blocked with picrotoxin (Figure 3D,E), indicating that I_h_ and Kir strongly influence synaptic integration even when inhibitory conductances are pharmacologically suppressed.

**Figure 3.**
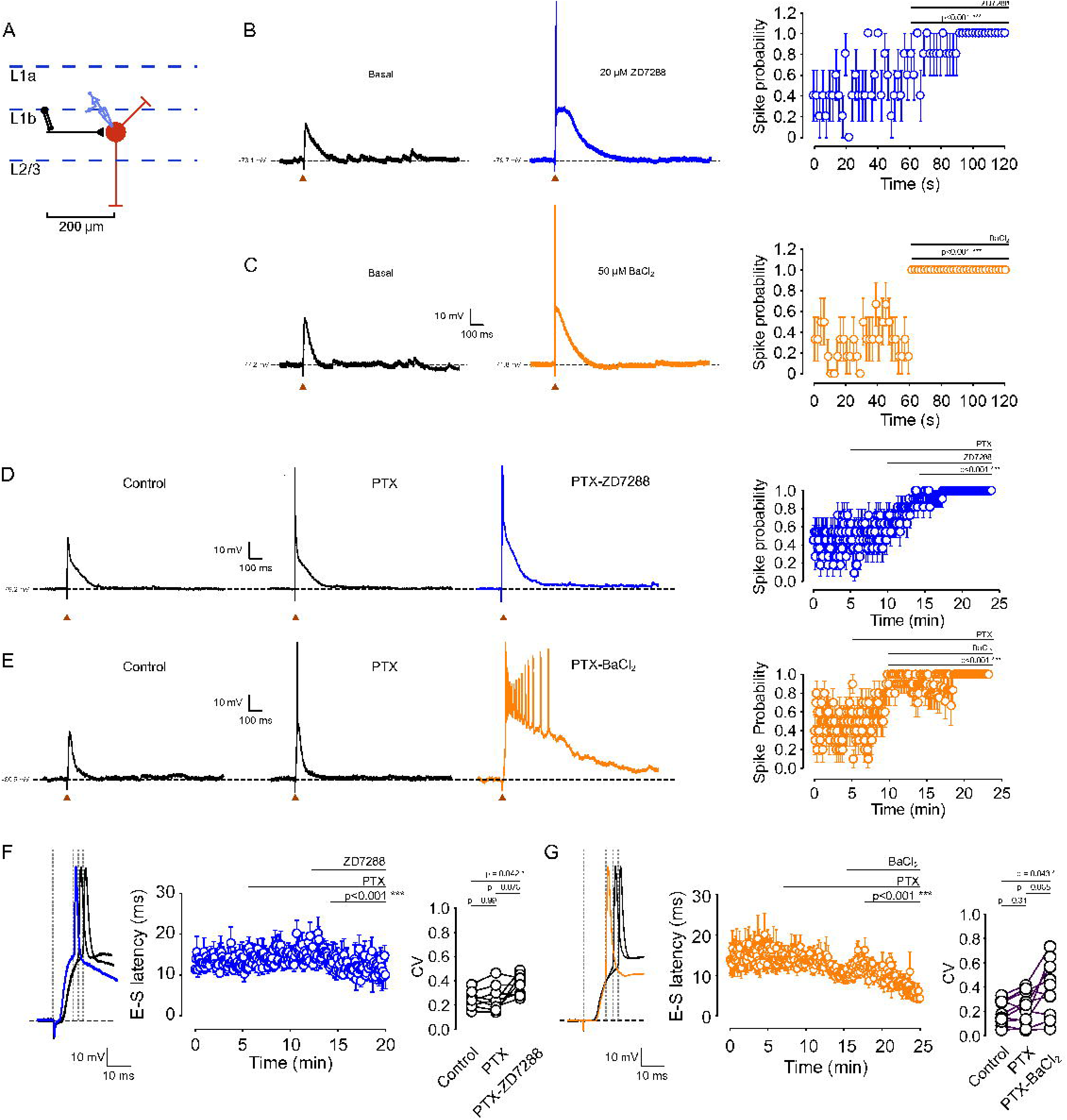
Inhibition of Kir and I_h_ currents increases spike probability in VIP interneurons. (A) Schematic of the recording configuration showing the distance between the stimulating electrode and the recorded neuron (200 µm), along with the pharmacological treatment protocol.(B) Left: Effect of 20 µM ZD-7288 on the evoked EPSP and spike probability induced by synaptic stimulation. Right: Summary plot of spike probability over time, before and after 20 µM ZD-7288 treatment (n = 5 cells/5 animals; p < 0.001).(C) Left: Effect of 50 µM BaCl_2_ on the evoked EPSP and spike probability induced by synaptic stimulation. Right: Summary plot of spike probability over time, before and after 50 µM BaCl_2_ treatment (n = 6 cells/6 animals; p < 0.001). (D) Left: Representative traces in control, 50 µM PTX, and 50 µM PTX + 20 µM ZD-7288. Right: Spike probability over time before and after PTX and PTX + ZD-7288 application (n = 8 cells/6 animals).(E) Left: Representative traces in control, 50 µM PTX, and 50 µM PTX + 50 µM BaCl_2_. Right: Spike probability over time before and after PTX and PTX + BaCl_2_ application (n = 10 cells/6 animals). (F) Left: Representative trace illustrating the EPSP-spike latency measurement. Right: Quantification of EPSP-spike latency over time in the presence of PTX and PTX + ZD-7288. (G) Left: Representative trace of spike latency. Right: Quantification of EPSP-spike latency over time in the presence of PTX and PTX + BaCl_2_. Data are presented as mean ± 95% C.I.

Moreover, we found that both ZD-7288 and BaCl_2_ increased the EPSP slope and decreased EPSP-spike latency (Figure 3F, G; Supplementary Figure 1), reflecting accelerated membrane depolarization toward threshold. Both drugs also reduced spike precision (increased latency variability), suggesting that I_h_ and Kir normally constrain the temporal window for EPSP-spike transformation. Together, these findings show that I_h_ and Kir exert potent control over EPSP integration and spike generation in L1b VIP interneurons, primarily by modifying subthreshold membrane properties rather than synaptic amplitude.

### 3.3. I_h_ and Kir differentially modulate intrinsic excitability of L1b VIP interneurons

We next assessed how I_h_ and Kir shape passive and active intrinsic properties. Blocking I_h_ with 20 µM ZD-7288 hyperpolarized the resting membrane potential (RMP) and increased input resistance, with no significant effect on membrane time constant or capacitance (Figure 4A; Supplementary Table 1). Hyperpolarizing current injections revealed that 20 µM ZD-7288 abolished the characteristic voltage sag, as expected for I_h_ inhibition. Despite increased R_in_, ZD-7288 produced no significant change in rheobase or firing frequency across the tested current range (Figure 4A), similarly, these effects were also observed at 5 µM ZD-7288 (Supplementary Figure 3, Supplementary Table 2), consistent with I_h_ playing its role in subthreshold integration.

**Figure 4.**
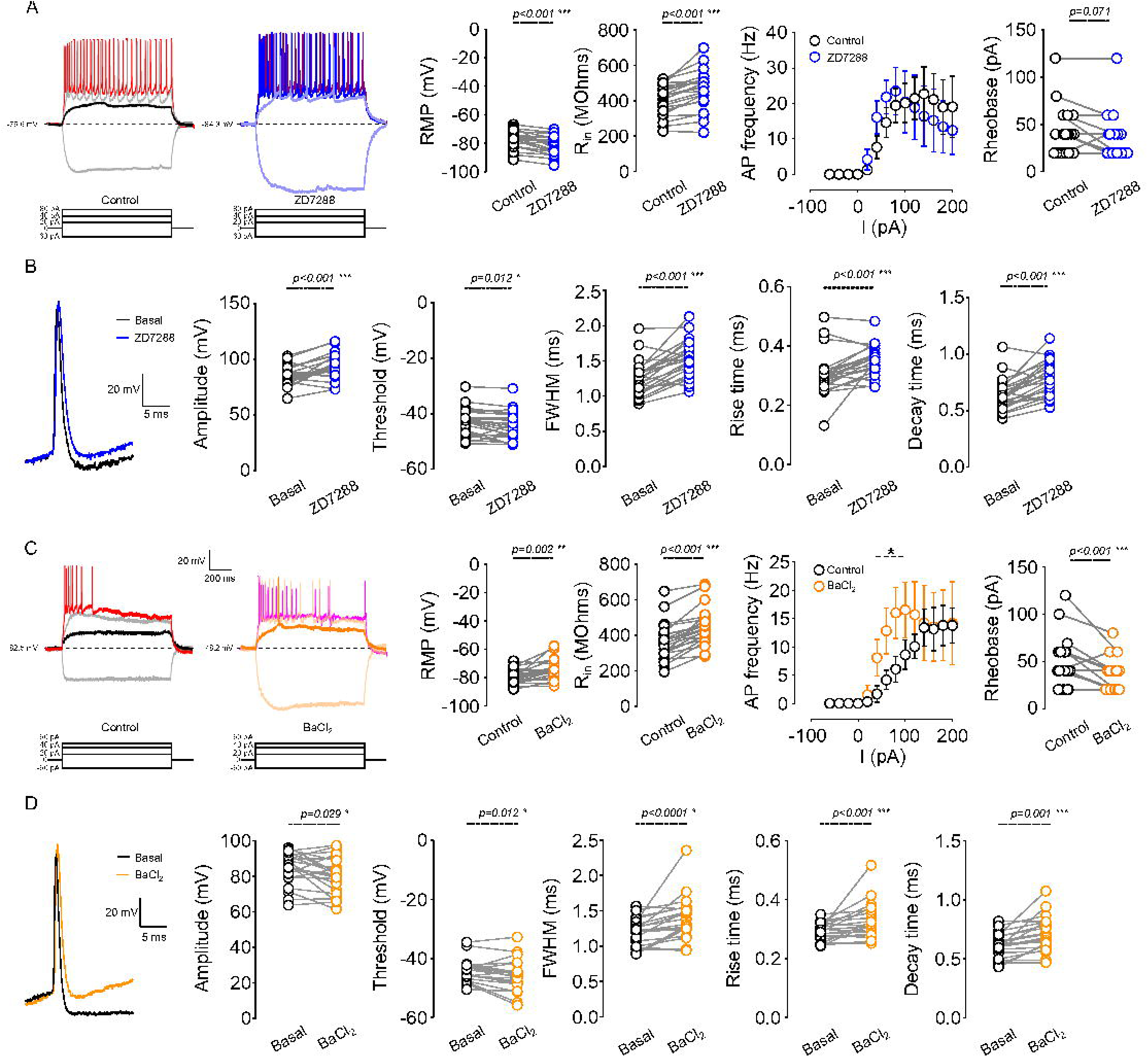
Inhibition of Kir and I_h_ conductances enhances the intrinsic excitability of layer 1b VIP interneurons. (A) Left: Representative traces showing the voltage response to current injections of -60 pA, 20, 40 pA, and 80 pA in mPFC L1 VIP interneurons in the presence of 20 µM ZD-7288. Right: Summary plots showing the effect of 20 µM ZD-7288 on resting membrane potential (RMP), input resistance (R_in_), AP frequency, and rheobase (n = 23 cells/18 animals). (B) Left: Representative traces showing the effect of ZD-7288 on the shape of the first action potential (AP) at rheobase. Right: Summary plots of the effect of ZD-7288 on AP amplitude, threshold, full width at half maximum (FWHM), rise time, and decay time (n = 23 cells/18 animals). (C) Left: Representative traces showing the effect of 50 µM BaCl_2_ on the voltage response evoked by -60 pA, 20, 40 pA, and 60 pA current injections. Right: Effect of 50 µM BaCl_2_ on RMP, R_in_, AP frequency, and rheobase (n = 22 cells/13 animals). (D) Left: Representative traces showing the effect of BaCl_2_ on AP shape. Right: Summary plots of the effect of BaCl_2_ on AP amplitude, threshold, FWHM, rise time, and decay time (n = 22 cells/13 animals). Data for AP firing frequency are shown as mean ± 95% C.I. Asterisks (*) indicate currents where p-values are statistically significant (p < 0.05). P-values are indicated above each graph.

Action-potential waveform analysis showed that ZD-7288 hyperpolarized the AP threshold, increased AP amplitude, and prolonged AP duration by slowing both rise and decay kinetics (Figure 4B; Supplementary Table 1), reflecting altered recruitment of voltage-gated Na^+^ and K^+^ channels at more hyperpolarized potentials. In contrast, blocking Kir with 50 µM BaCl_2_ depolarized the RMP, increased R_in_, and prolonged the membrane time constant while leaving capacitance unchanged (Figure 4C; Supplementary Table 1). Hyperpolarizing steps produced larger voltage deflections but also revealed an increase in sag amplitude, suggesting recruitment of I_h_. Current-step protocols demonstrated that BaCl_2_ substantially decreased rheobase and increased firing frequency at moderate current injections (40-100 pA), although this facilitation plateaued at higher current amplitudes (Figure 4E). As with I_h_ inhibition, BaCl_2_ hyperpolarized AP threshold, increased AP amplitude, and prolonged AP duration (Figure 4D). Together, these findings indicate that I_h_ and Kir are both active at rest and jointly regulate passive properties and action-potential generation, but exert distinct influences on firing gain and excitability: I_h_ primarily shapes subthreshold responsiveness, whereas Kir sets the depolarization threshold for AP initiation and determines gain in the low-current regime.

### 3.4. Kir inhibition unmasks an I_h_-dependent voltage sag in L1b VIP interneurons

Because BaCl_2_ increased R_in_ and hyperpolarization amplitude, we hypothesized that Kir inhibition enhances sag amplitude by increasing I_h_ recruitment. We quantified sag during a -60 pA current step before and after BaCl_2_, and again after BaCl_2_ + ZD-7288 (Figure 5A). In 85% of neurons (11/13), BaCl_2_ significantly increased sag amplitude (Figure 5B,C). Subsequent application of ZD-7288 abolished this enhanced sag, demonstrating that the BaCl_2_-induced sag is I_h_-dependent. These results indicate that under baseline conditions constitutive Kir activity stabilizes the membrane potential in a range that limits I_h_ activation during modest hyperpolarizations, effectively masking sag. Removal of Kir shifts the operating point into a regime where I_h_ becomes strongly engaged, unmasking the sag response. This interaction suggests a dynamic balance between Kir and I_h_ in shaping subthreshold voltage trajectories.

**Figure 5.**
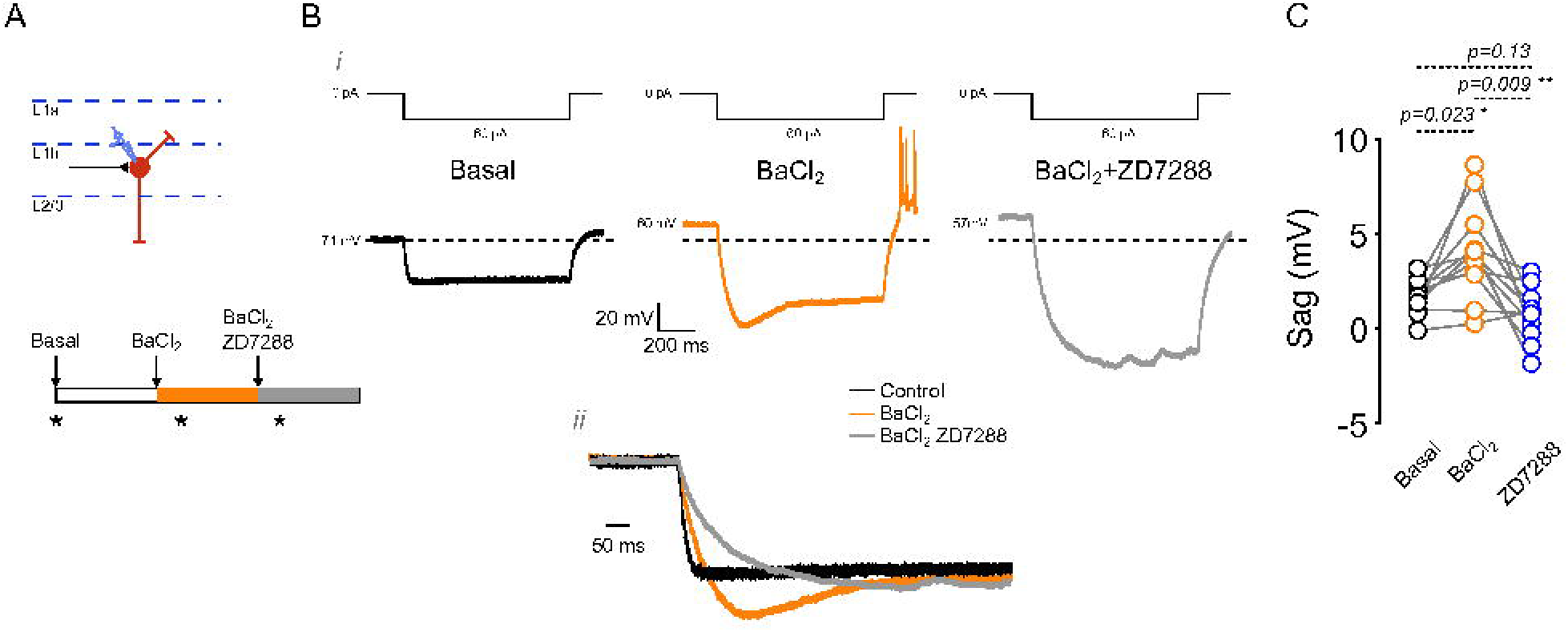
Kir inhibition unmasks a ZD-7288-sensitive voltage sag. (A) Schematic representation of the recording area and the pharmacological treatment protocol. (B) (i) Representative traces showing the response to a -60 pA somatic current injection in control, 50 µM BaCl_2_, and 50 µM BaCl_2_ + 20 µM ZD-7288 conditions. The current injection protocol is shown above the traces. (ii) Higher magnification of the boxed area showing the effect of treatments on sag amplitude. (C) Summary plot quantifying the effect of BaCl_2_ and BaCl_2_ + ZD-7288 on sag amplitude (n = 11 cells/11 animals). P-values are shown above the graph.

### 3.5. Voltage-clamp identification of I_h_ and Kir currents and HCN channel expression in L1b VIP interneurons

To directly measure subthreshold currents, we performed voltage-clamp recordings from -130 to -50 mV using pharmacological isolation (Figure 6). Under control conditions, L1b VIP interneurons displayed a small inward current with an apparent reversal around -82 mV (uncorrected for the liquid junction potential; Figure 6A). Application of ZD-7288 reduced inward current between -130 and -100 mV, yielding a ZD-sensitive component of ∼-18 pA at -130 mV with an apparent reversal near -76 mV (Figure 6B), consistent with an I_h_-like mixed cation current. Application of BaCl_2_ revealed a Ba^2+^-sensitive inwardly rectifying current of ∼-30 pA at -130 mV with reversal around -82 mV (Figure 6C, D), consistent with Kir activation. Although these currents are small in amplitude, they operate on a high-resistance membrane, indicating that even tens of picoamps can substantially shape excitability. To determine the molecular identity of I_h_ channels, we performed immunofluorescence in VIP^Cre^;Ai9 mice. VIP neurons were confined to L1b and exhibited predominantly bipolar morphology (Figure 7). HCN1 immunoreactivity was weak and primarily somatic, and not all VIP neurons expressed detectable HCN1. No HCN2 signal was observed in L1b or L2/3 VIP interneurons (Supplementary Figure 2). In line with previous reports, we identified HCN1-labeled distal dendrites originating from deep layers. This suggests that the low expression of HCN1 observed in VIP interneurons is cell-type specific rather than a general labeling deficit. Furthermore, HCN1 expression in the prefrontal cortex followed the classical gradient described by Lörincz et al. [34], with density increasing toward the distal dendrites in pyramidal neurons (Figure 7, red arrows, Supplementary Figure 4). These findings support the presence of a small somatic I_h_ conductance mediated mainly by HCN1, consistent with the modest I_h_ amplitude observed in voltage clamp.

**Figure 6.**
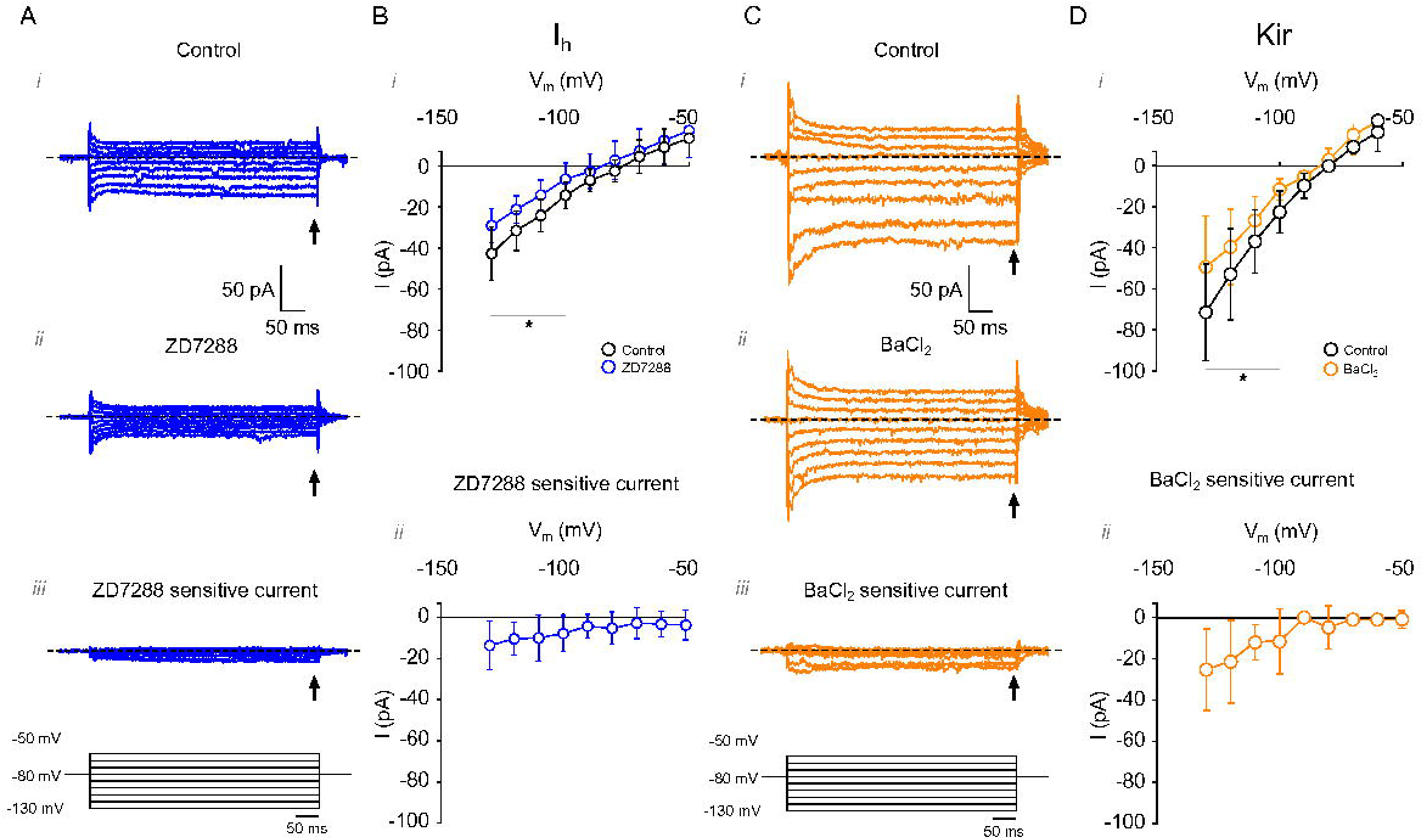
I_h_ and Kir currents in mPFC L1 VIP interneurons. (A) Representative current traces measured using a voltage step protocol from -130 to -50 mV: (i) before, (ii) after ZD-7288 application, and (iii) the ZD-7288-sensitive difference current (Control - ZD-7288). (B) Quantification of (i) total current and (ii) ZD-7288-sensitive current (n = 6 cells/6 animals). (C) Representative current traces measured using a voltage step protocol from -130 to -50 mV: (i) before, (ii) after BaCl_2_ application, and (iii) the BaCl_2_-sensitive difference current (Control - BaCl_2_). (D) Quantification of (i) total current and (ii) BaCl_2_-sensitive current (n = 6 cells/6 animals). Data are presented as mean ± 95% C.I. Arrows indicate the time point where measurements were taken. *p < 0.05.

**Figure 7.**
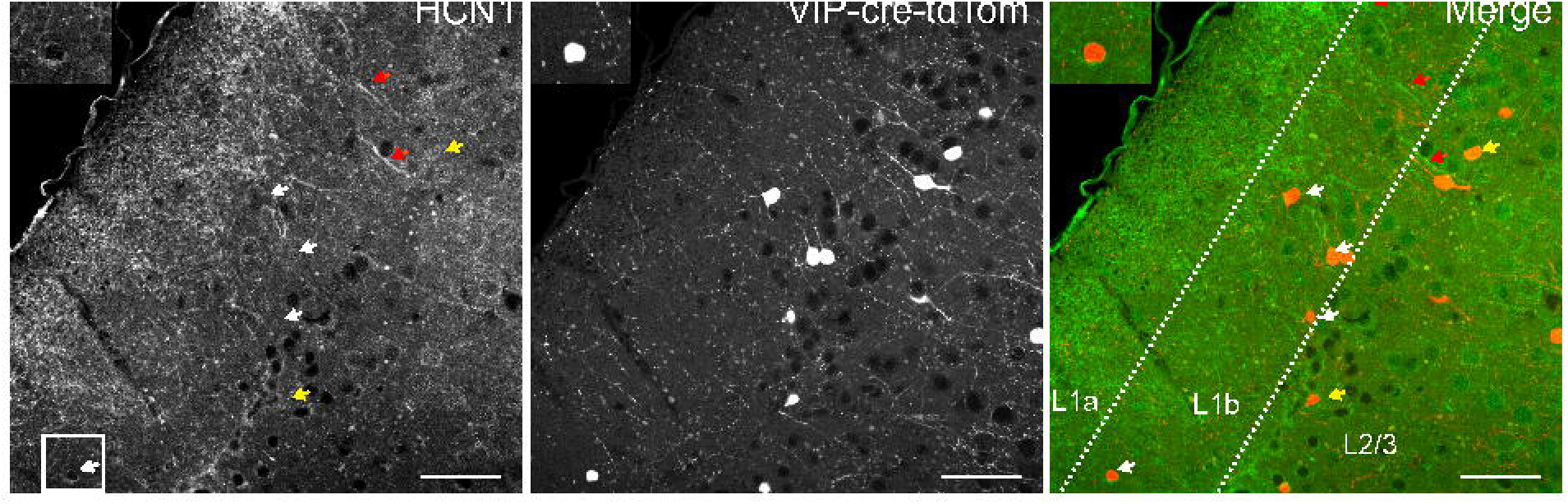
HCN1 expression in mPFC L1 VIP interneurons. Representative immunofluorescence images showing labeling for HCN1 (green), tdTomato (red), and the merged signal. Lines indicate layer boundaries. Inset shows a higher magnification view of the cells in the enclosed square. Scale bar = 50 µm. White arrow indicates neurons in layer 1; yellow arrow indicates VIP interneurons in layer 2; red arrows indicate HCN1 expression in dendrites coming from deep layers.

## 4. DISCUSSION

In this study, we show that the subthreshold currents I_h_ and Kir are constitutively active in L1b VIP interneurons of the medial prefrontal cortex and critically regulate their passive properties, intrinsic excitability, and EPSP-spike coupling. By combining whole-cell recordings, synaptic stimulation, voltage-clamp isolation of subthreshold currents, and immunofluorescence, we demonstrate that these two conductances jointly shape how VIP interneurons integrate synaptic inputs and determine their responsiveness to excitatory drive. Our findings position I_h_ and Kir as key regulators of signal filtering in layer 1, where VIP neurons function as important modulators of disinhibition and top-down information flow across cortical layers.

### 4.1 I_h_ conductance regulates subthreshold integration and EPSP-spike coupling

Our results demonstrate that I_h_ is active near the resting membrane potential of L1b VIP interneurons, where it acts as a depolarizing conductance that lowers input resistance and constrains EPSP-spike coupling. Pharmacological inhibition of I_h_ with ZD-7288 hyperpolarized the resting membrane potential, increased input resistance, abolished the voltage sag, and enhanced both the probability and temporal precision with which excitatory inputs triggered action potentials. These effects closely mirror those reported in CA1 pyramidal neurons, where I_h_ shapes dendritic depolarization, synaptic integration, and long-term plasticity [15,23,34], supporting a conserved role for I_h_ in regulating subthreshold excitability.

Notably, I_h_ inhibition did not alter spontaneous EPSP or miniature EPSC amplitude or frequency, indicating that I_h_ does not significantly influence presynaptic glutamate release or postsynaptic glutamate receptor responsiveness under basal conditions. Instead, its primary role appears to be the regulation of intrinsic membrane responsiveness, consistent with observations across multiple neuronal classes [15,18,19]. Voltage-clamp recordings revealed a small ZD-7288-sensitive inward current (∼-18 pA at -130 mV), and immunofluorescence analysis showed weak, predominantly somatic HCN1 expression with no detectable HCN2 signal in L1b VIP interneurons. These findings align with reports of relatively modest I_h_ amplitudes in interneurons [35,36]. Despite its small magnitude, I_h_ exerted substantial functional effects, likely due to the high input resistance characteristic of L1 interneurons [30], which amplifies the voltage impact of small conductances.

Given the high input resistance of L1 VIP interneurons, even small I_h_ active near rest can produce millivolt-scale shifts in membrane potential (ΔV ≈ I × R_in_), consistent with our observed changes in RMP and EPSP-spike coupling. In addition, weak somatic HCN1 immunoreactivity may underestimate functionally relevant channel pools in fine dendrites or other subcellular compartments. Importantly, the main intrinsic effects were replicated at 5 µM ZD-7288 (Supplementary Figure 3), supporting the interpretation that they largely reflect I_h_ blockade rather than high-dose off-target actions.

The enhancement of EPSP-spike coupling following I_h_ inhibition can therefore be explained by an increase in input resistance, which promotes EPSP summation and facilitates recruitment of conductances such as NMDARs. The absence of changes in sEPSC and mEPSC properties further supports a predominantly postsynaptic mechanism. Although ZD-7288 has been reported to affect T-type Ca^2+^ and Na^+^ channels at higher concentrations [37,38], our analyses focused on subthreshold membrane potentials where these channels are minimally active, supporting the interpretation that the observed effects primarily reflect I_h_ blockade. Nevertheless, because high concentrations of ZD-7288 can enhance presynaptic glutamate release in certain circuits [39,40], subtle presynaptic contributions cannot be completely excluded.

Together, these data indicate that I_h_ functions as a stabilizing, depolarizing leak conductance in L1b VIP interneurons that narrows the temporal window for synaptic integration and limits spike probability. By doing so, I_h_ fine-tunes the gain and timing of excitatory input-output coupling in these cells, positioning it as a key regulator of excitability and temporal precision in layer 1 cortical microcircuits.

### 4.2. Kir conductance shapes resting potential, excitability, and the temporal integration window

Kir inhibition with BaCl_2_ produced effects opposite to those of I_h_ inhibition: depolarization of RMP, increased R_in_ and membrane time constant, reduced rheobase, and increased firing frequency for low to moderate current injections. These findings demonstrate that Kir channels normally provide a stabilizing conductance that constrains excitability, consistent with the canonical role of inwardly rectifying potassium channels [16,21]. Voltage-clamp recordings revealed a Ba^2+^-sensitive inwardly rectifying current (∼-30 pA at -130 mV), with characteristics similar to Kir2.x channels. Although scRNAseq datasets indicate that GIRK (Kir3.x) subunits are expressed in VIP interneurons [41–43], the presence of a constitutively active Ba^2+^-sensitive current suggests additional Kir family members may contribute. Similar interactions among HCN, Kir2.x, and K-leak channels have been described as key determinants of dendritic excitability in frontal cortex pyramidal neurons [21]. Kir inhibition enhanced EPSP-spike coupling and broadened EPSPs, consistent with increased R_in_ and prolonged membrane integration time. Prior studies showed that Kir inhibition increases EPSP amplitude and temporal summation in striatal and cortical neurons [21,43]. Our data extend this principle to L1b VIP interneurons, showing that Kir channels exert tonic control over synaptic integration by limiting depolarization and narrowing the integration window.

Beyond shifting the resting membrane potential, Kir activation reduces VIP interneuron excitability through shunting inhibition. By increasing total membrane conductance, Kir channels decrease R_in_, so identical EPSCs produce smaller voltage deflections (EPSPs), effectively filtering weak or subthreshold inputs even independent of any hyperpolarizing shift in membrane potential. This shunting mechanism is particularly relevant in VIP interneurons, where high basal R_in_ makes excitability highly sensitive to modest changes in ionic conductance.

### 4.3. Interplay between I_h_ and Kir establishes a dynamic balance governing voltage trajectories and spike timing

Our results indicate that Kir inhibition unmasks an I_h_-dependent voltage sag. Under baseline conditions, hyperpolarizing current injections produced minimal sag, suggesting limited I_h_ recruitment. However, blocking Kir strongly increased sag amplitude, and subsequent ZD-7288 application abolished this effect. This indicates that Kir current shunts I_h_ current and stabilizes the membrane potential, effectively masking sag.

This interplay resembles observations in CA1 pyramidal neurons, where GIRK activation reduces I_h_ recruitment by preventing hyperpolarizing shifts in membrane potential[44]. Contrary, in L1b VIP interneurons, removing Kir reduces current shunting thus unmasking I_h_ current, increasing sag amplitude and altering voltage trajectories. We also found that blocking GABA-A receptors alone had little effect on spontaneous synaptic activity or RMP in L1b VIP interneurons, consistent with reports that L1 receives relatively sparse local inhibition [1,30]. However, during evoked excitation, picrotoxin increased spike probability, and this effect was greatly amplified when Kir was inhibited, suggesting that Kir-mediated shunting normally masks the influence of feedforward inhibition. Removal of Kir thus increases membrane gain and enhances the impact of both excitatory and inhibitory synaptic inputs.

In this work, we utilized mice ranging from 1 to 3 months of age, a period spanning late adolescence to young adulthood. While previous literature suggests that Layer 1 interneuron markers are expressed as early as P14, it is critical to acknowledge that the prefrontal cortex (PFC) undergoes a protracted maturation process [45]. Sexual maturity in mice typically occurs around 6 weeks, and the refinement of PFC connectivity and synaptic pruning continues throughout the 1-3 month window [46,47]. We did not observe clear age-related trends within this range, but the study was not powered to detect subtle developmental effects.

Together, these results show that I_h_ and Kir are active near the resting membrane potential of mPFC L1b VIP interneurons, where they jointly regulate the voltage range over which synaptic inputs are integrated. By shaping subthreshold filtering, intrinsic excitability, and EPSP-spike coupling, the interplay between these conductances establishes a dynamic balance that determines the timing, probability, and precision of spike generation. This coordinated regulation positions VIP interneurons as finely tunable modulators of disinhibition and information flow within the cortical column.

## Ethics declaration

All experiments were conducted according to animal protocols approved by the Ethics Committee of the Universidad de Santiago de Chile (N° 193/2022), according to the rules and guidelines of the National Research and Development Agency (ANID)

## Availability of data and materials

The data supporting this study’s findings are available from the corresponding author upon request.

## Acknowledgments

The authors also thank the Microscopy Core Facility at Universidad de Santiago de Chile, funded by Programa de Equipamiento Científico y Tecnológico (FONDEQUIP) EQM150069.

## Author Contributions

E.L-S and C.M. conceived the experiments and developed the analyses. C.M., D.R., C.C, P.L. and EL-S. performed the experiments and analyzed the data. E.L-S wrote the manuscript, and all authors edited the manuscript.

## 6. FIGURES

**Supplementary Table 1**

Electrophysiological characteristics of VIP interneurons before and after 20 µM ZD-7288 and 50 µM BaCl_2_ treatment.

**Supplementary Table 2**

Electrophysiological characteristics of VIP interneurons before and after 5 µM ZD-7288 treatment.

**Supplementary Figure 1**

EPSP input/output curves of mPFC L1b VIP interneurons. Input/output (I/O) curves recorded from L1b VIP interneurons following stimulation in L1b, 200 µm from the soma. (A) Slope of the I/O relationship in the presence of 20 µM ZD-7288. (B) EPSP amplitude I/O relationship in the presence of 20 µM ZD-7288 (n = 5 cells/5 animals). (C) Slope of the I/O relationship in the presence of 50 µM BaCl_2_. (D) EPSP amplitude I/O relationship in the presence of 50 µM BaCl_2_ (n = 6 cells/6 animals).

**Supplementary Figure 2**

(A) Representative immunofluorescence images showing labeling for HCN2, tdTomato, and the merged signal. (B) Representative immunofluorescence images showing the negative control for the primary antibody, tdTomato, and the merged signal. Lines indicate layer boundaries. Inset shows a higher magnification view of the cells in the enclosed square. Scale bar = 50 µm.

**Supplementary Figure 3**

Effect of 5 µM ZD-7288 on layer 1 VIP interneurons in the mPFC. (A) (i) Representative voltage responses to somatic current injections of -60, 20, and 40 pA. Summary graphs show (ii) resting membrane potential, (iii) input resistance, (iv) sag, (v) action potential firing frequency, and (vi) rheobase. (vii) Representative voltage responses to the same current injections, with summary graphs of (viii) action potential threshold, (ix) amplitude, (x) full width at half maximum, (xi) rise time, and (xii) decay time (n = 3). (B) Representative traces illustrating the effects of 5 µM ZD-7288 on sEPSP amplitude and frequency (right panels) (n = 4). (C) Summary graphs showing the effects of 5 µM ZD-7288 on mEPSC amplitude and frequency. (D) Representative traces showing the effects of 5 µM ZD-7288 on eEPSPs under (i) control conditions, (ii) picrotoxin, and (iii) picrotoxin + 5 µM ZD-7288. (iv) Summary graph of spike probability in the presence of 5 µM ZD-7288 (n = 2).

**Supplementary Figure 4**

HCN1 expression in the mPFC and hippocampal CA1 at P35. (A) Representative images of HCN1 (green) and MAP2 (red) immunolabeling in mPFC layers 1 and 2/3, highlighting HCN1 signal in dendritic processes (yellow arrows). (B) Representative images of HCN1 (green) and MAP2 (red) immunolabeling in the hippocampal CA1 region. Scale bar, 20 µm.

