## Supplementary figures and images for "Subthreshold Kir and I_h_ currents modulate excitability of Layer 1 VIP interneurons in the medial prefrontal cortex"

### Supplemental figure 1

#### Supplementary Figure 1

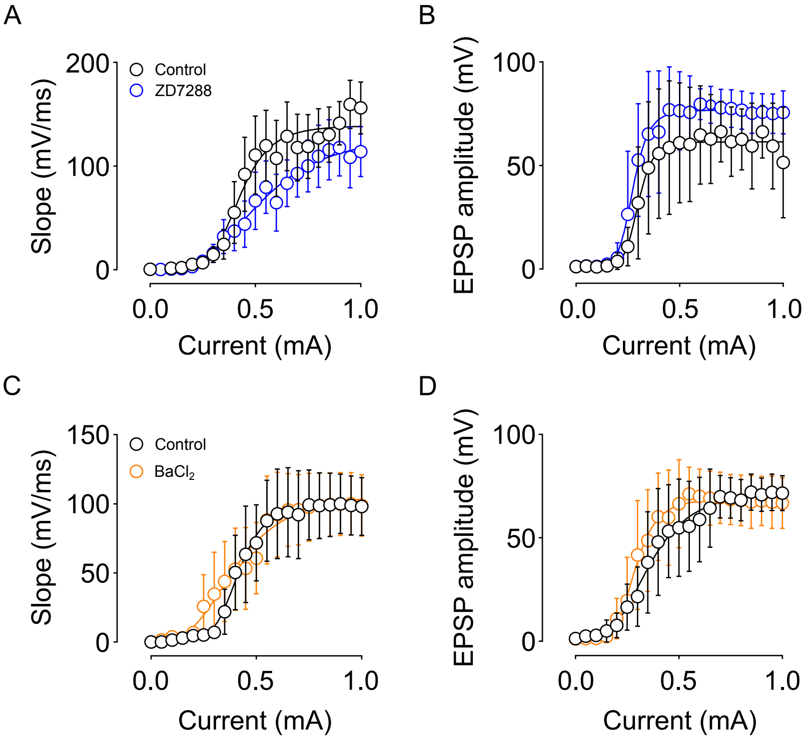

### Supplementary figure 2

Supplementary Figure 2

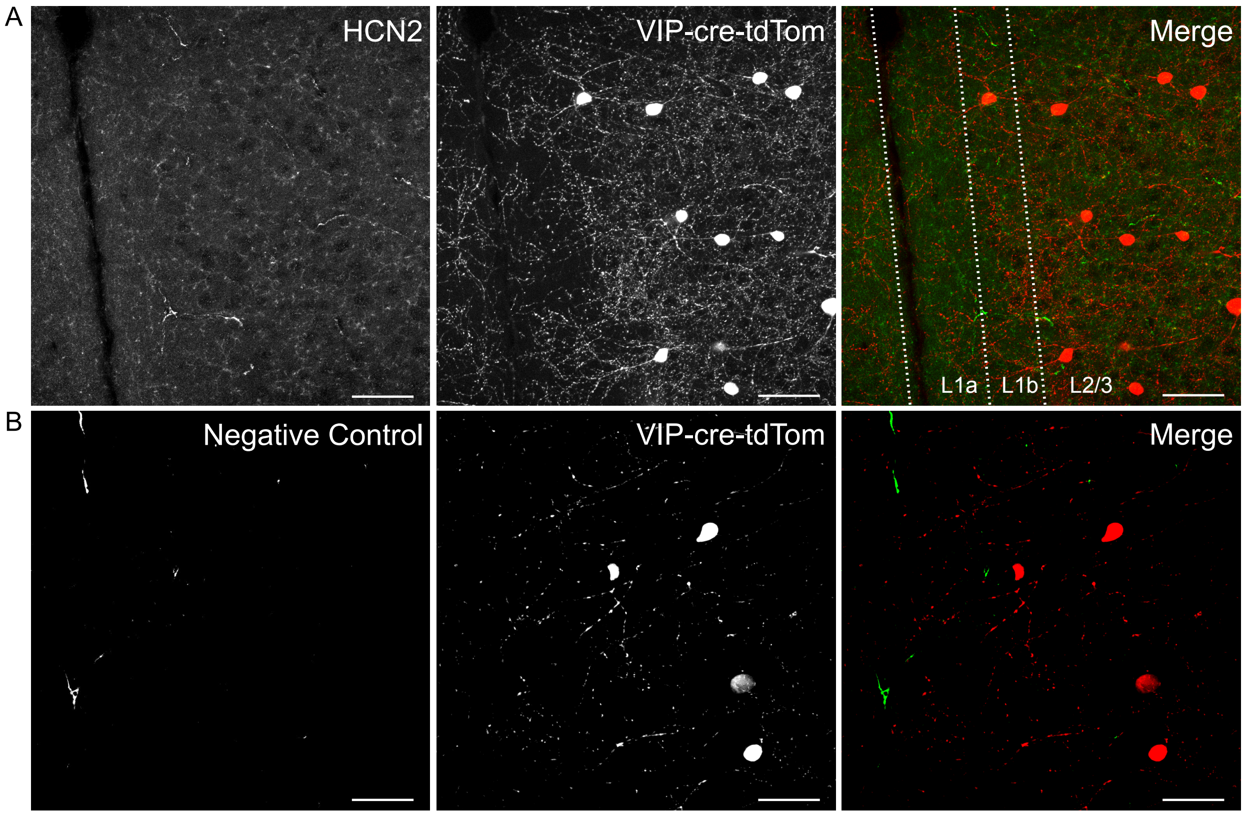

### Supplementary figure 3

#### Supplementary figure 3

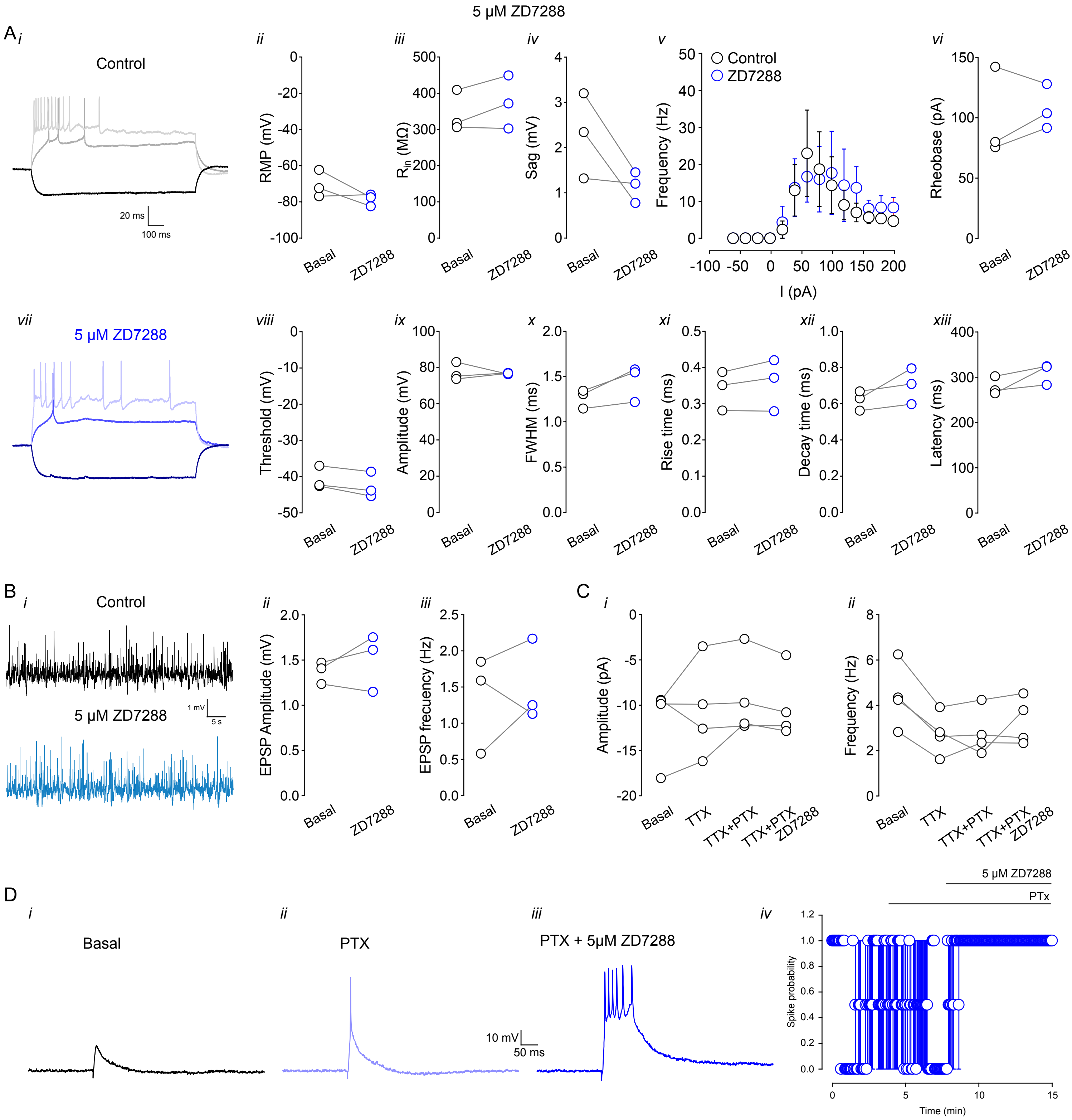

### Supplementary figure 4

Supplementary Figure 4

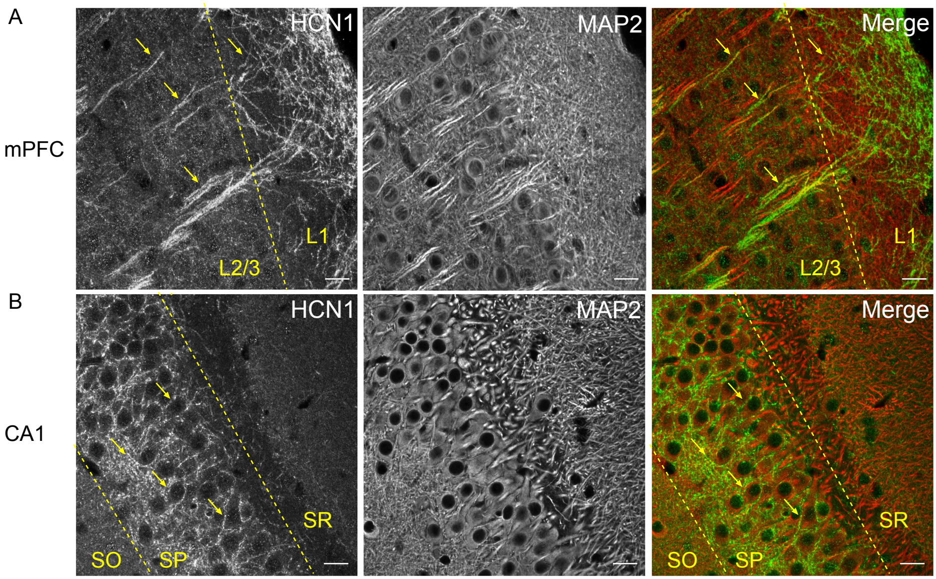
