## Supplementary Table 1 for "Subthreshold Kir and I_h_ currents modulate excitability of Layer 1 VIP interneurons in the medial prefrontal cortex"

### Action potential parameters

| | Control | | ZD7288 | | | | | Control | | $BaCl_2$ | | | | |
|---|---|---|---|---|---|---|---|---|---|---|---|---|---|---|
| | ***Mean*** | ***SD*** | ***Mean*** | ***SD*** | ***Statistical test*** | ***p-value*** | | ***Mean*** | ***SD*** | ***Mean*** | ***SD*** | ***Statistical test*** | ***p-value*** | |
| **n** | N = 18, n = 23 | | | | | | | N = 13, n = 22 | | | | | | |
| **RMP (mV)** | -77.2 | 6.2 | -81.5 | 6.4 | Paired t test | 0.0001 | * | -79.1 | 5.1 | -74.7 | 7.2 | Paired t test | 0.002 | * |
| **Rin (MOhms)** | 410.8 | 87.9 | 468.2 | 110.8 | Paired t test | 0.0001 | * | 362.9 | 105.5 | 446.3 | 105.2 | Paired t test | 0.0001 | * |
| **Cm (pF)** | 55.6 | 13.4 | 50.5 | 17.2 | Paired t test | 0.115 | | 50.6 | 15.6 | 59.7 | 18.8 | Paired t test | 0.0879 | |
| **Tau (ms)** | 22.9 | 7.8 | 23.6 | 16.9 | Wilcoxon matched-pairs signed rank test | 0.2861 | | 18.4 | 7.0 | 35.6 | 12.8 | Wilcoxon matched-pairs signed rank test | 0.0001 | * |
| **Rheobase (pA)** | 38.3 | 24.8 | 33.1 | 23.0 | Wilcoxon matched-pairs signed rank test | 0.071 | | 47.7 | 25.4 | 33.1 | 16.8 | Wilcoxon matched-pairs signed rank test | 0.002 | * |
| **Threshold (mV)** | -42.9 | 5.1 | -44.3 | 5.2 | Paired t test | 0.0122 | * | -44.4 | 3.9 | -46.0 | 5.5 | Paired t test | 0.0124 | * |
| **Amplitude (mV)** | 85.3 | 8.9 | 93.8 | 11.4 | Paired t test | 0.0004 | * | 84.5 | 9.1 | 80.7 | 10.3 | Paired t test | 0.029 | * |
| **Half time (ms)** | 1.3 | 0.2 | 1.5 | 0.3 | Paired t test | 0.0001 | * | 1.2 | 0.2 | 1.4 | 0.3 | Wilcoxon matched-pairs signed rank test | 0.0001 | * |
| **Rise (ms)** | 0.3 | 0.1 | 0.4 | 0.1 | Wilcoxon matched-pairs signed rank test | 0.0002 | * | 0.3 | 0.0 | 0.3 | 0.1 | Wilcoxon matched-pairs signed rank test | 0.0003 | * |
| **Decay (ms)** | 0.6 | 0.1 | 0.8 | 0.2 | Paired t test | 0.0001 | * | 0.6 | 0.1 | 0.7 | 0.1 | Paired t test | 0.0003 | * |
| **AHP (ms)** | 4.9 | 6.4 | 2.2 | 5.7 | Paired t test | 0.0565 | | -4.1 | 7.2 | -1.1 | 6.9 | Paired t test | 0.1332 | |
| **Latency (ms)** | 280.0 | 42.5 | 261.3 | 32.3 | Wilcoxon matched-pairs signed rank test | 0.0043 | * | 281.1 | 51.1 | 242.7 | 32.7 | Wilcoxon matched-pairs signed rank test | 0.0003 | * |
| **Sag (mV)** | 3.4 | 1.8 | 1.0 | 1.2 | Wilcoxon matched-pairs signed rank test | 0.0001 | * | 1.7 | 0.9 | 4.1 | 2.5 | Tukey's multiple comparisons test | 0.0233 | * |

**** Statistical significance***

### sEPSP parameters

| | Control | | PTX | | PTX-ZD7288 | | | | Control | | PTX | | $PTX-BaCl_2$ | | | |
|---|---|---|---|---|---|---|---|---|---|---|---|---|---|---|---|---|
| **n** | N = 4, n = 4 | | | | | | | | N = 4, n = 6 | | | | | | | |
| | ***Mean*** | ***SD*** | ***Mean*** | ***SD*** | ***Mean*** | ***SD*** | ***Statistical test*** | ***p-value*** | ***Mean*** | ***SD*** | ***Mean*** | ***SD*** | ***Mean*** | ***SD*** | ***Statistical test*** | ***p-value*** |
| **EPSP amplitude (mV)** | 1.43 | 0.548 | 1.453 | 0.5923 | 1.676 | 0.8434 | one-way Anova | 0.376 | 1.3 | 0.35 | 1.4 | 0.47 | 1.5 | 0.51 | one-way Anova | 0.2837 |
| **EPSP frequency (Hz)** | 0.91 | 0.5448 | 0.78 | 0.3803 | 1.13 | 0.3479 | one-way Anova | 0.2749 | 1.6 | 1.5 | 1.1 | 1.6 | 1.3 | 1.3 | one-way Anova | 0.0665 |

### EPSC parameters

| | Control | | TTX | | TTX-PTX | | TTX-PTX-ZD7288 | | | | Control | | TTX | | TTX-PTX | | $PTX-BaCl_2$ | | | |
|---|---|---|---|---|---|---|---|---|---|---|---|---|---|---|---|---|---|---|---|---|
| **n** | N = 5, n = 5 | | | | | | | | | | N = 5, n =5 | | | | | | | | | |
| | ***Mean*** | ***SD*** | ***Mean*** | ***SD*** | ***Mean*** | ***SD*** | ***Mean*** | ***SD*** | ***Statistical test*** | ***p-value*** | ***Mean*** | ***SD*** | ***Mean*** | ***SD*** | ***Mean*** | ***SD*** | ***Mean*** | ***SD*** | ***Statistical test*** | ***p-value*** |
| **EPSC amplitude (pA)** | -12.31 | 2.221 | -11.92 | 2.221 | -12.92 | 2.391 | -12.86 | 2.76 | one-way Anova | 0.2614 | -10.5 | 3.16 | -8.366 | 2.79 | -7.936 | 1.59 | -8.166 | 2.053 | one-way Anova | 0.0936 |
| **EPSC frequency (Hz)** | 2.222 | 2.472 | 1.93 | 1.918 | 1.908 | 1.954 | 2.13 | 2.237 | one-way Anova | 0.1495 | 1.793 | 0.9125 | 1.49 | 0.8068 | 1.572 | 0.6445 | 1.594 | 0.9467 | one-way Anova | 0.1495 |
