## Supplementary table 2 for "Subthreshold Kir and I_h_ currents modulate excitability of Layer 1 VIP interneurons in the medial prefrontal cortex"

### Supplementary table 2. Parameter for 5 μM ZD-7288

| *Excitability* | Control | | ZD7288 | |
|---|---|---|---|---|
| N | 3 | | | |
| | *Mean* | *SD* | *Mean* | *SD* |
| RMP (mV) | -70.5 | 7.5 | -78.6 | 3.4 |
| Rin (Mohms) | 344.6 | 56.2 | 374.7 | 73.4 |
| Cm (pF) | 76.2 | 27.3 | 60.5 | 10.5 |
| Tau (ms) | 30.2 | 7.9 | 24.1 | 3.1 |
| Rheobase (pA) | 99.4 | 37.2 | 107.9 | 18.5 |
| Threshold (mV) | -40.6 | 3.2 | -42.6 | 3.5 |
| Amplitude (mV) | 77.3 | 5.0 | 76.7 | 0.4 |
| Half time (ms) | 1.3 | 0.1 | 1.4 | 0.2 |
| Rise time (ms) | 0.3 | 0.1 | 0.4 | 0.1 |
| Decay time (ms) | 0.6 | 0.1 | 0.7 | 0.1 |
| Latency (ms) | 279.7 | 20.2 | 310.5 | 23.5 |
| Sag (mV) | 2.3 | 0.9 | 1.1 | 0.3 |

| *sEPSP* | Control | | ZD7288 | |
|---|---|---|---|---|
| N | 3 | | | |
| | *Mean* | *SD* | *Mean* | *SD* |
| Amplitude (mV) | 1.37 | 0.12 | 1.51 | 0.32 |
| Frequency (Hz) | 1.34 | 0.67 | 1.52 | 0.57 |

| *mEPSC* | Control | | TTx | | TTX-PTX | | TTX-PTX-ZD7288 | |
|---|---|---|---|---|---|---|---|---|
| N | 4 | | | | | | | |
| | *Mean* | *SD* | *Mean* | *SD* | *Mean* | *SD* | *Mean* | *SD* |
| Amplitude (pA) | -11.7 | 4.2 | -10.5 | 5.4 | -9.2 | 4.5 | -10.1 | 3.8 |
| Frequency (Hz) | 4.4 | 1.4 | 2.7 | 0.9 | 2.8 | 1.0 | 3.3 | 1.0 |
